# Distinct talkers combat catastrophic failures of spatial attention due to interruption

**DOI:** 10.64898/2025.12.16.694683

**Authors:** Wusheng Liang, Abigail L. Noyce, Christopher A. Brown, Barbara G. Shinn-Cunningham

## Abstract

Task-irrelevant features can impact formation of auditory objects and influence the effectiveness of selective attention, including the buildup of attention over time. Using a previously established paradigm exploring the effects of random interruptions on spatial selective attention, this study explores how the task-irrelevant feature of talker identity impacts the buildup of spatial attention and whether it alters the impact of interruptions. Participants performed a sequence recall task in which participants were presented with two competing syllable sequences coming from different spatial directions and were asked to report the syllable sequence coming from the target direction. On half of the trials, an unpredictable, novel interrupting sound occurred, disrupting attentional focus. Two experiments explored how talker influenced performance, specifically, whether 1) making the two streams come from different talkers facilitates task performance and reduces the impact of interruption compared to when the streams are spoken by the same talker, and 2) talker discontinuity interferes with attention buildup and harms syllable recall performance compared to when the talker is the same from one syllable to the next. Our results showed that distinct talker features, though task-irrelevant in this spatial task, significantly improved syllable recall performance and reduced the impact of interrupters. Further, irrelevant talker discontinuities damaged attention buildup and reduced syllable recall performance.

## I. Introduction

When multiple sound sources produce signals simultaneously, a listener needs to segregate the sources and focus attention on the one of interest in order to understand it (Cherry, 1953). Over the last seventy years, many studies have explored the mechanisms by which listeners segregate and select sound sources to solve this “cocktail party problem.” (Bee & Micheyl, 2008; Bronkhorst, 2015; Qian et al., 2018). But there is another, equally common but less well-studied problem in everyday auditory environments: there is always a chance of interruption—unpredictable sound events may grab our attention involuntarily, through automatic bottom-up processes (Huang & Elhilali, 2020; Zhao et al., 2019). The interplay between top-down selective attention and bottom-up interruption helps balance our constant need to be able to focus on known sounds while staying alert to new events.

This study builds on previous online experiments examining factors that influence the interaction between top-down spatial attention and bottom-up interruptions (Liang et al., 2022, 2025). The current study builds on these earlier efforts by focusing on how interruptions interact with task-irrelevant acoustic features of competing streams during spatial selective attention.

In complex acoustic environments, the spectrotemporal structure of natural, independent sounds typically provides sufficient information for the auditory system to segregate different sources, even when they are acoustically similar—for example, two different utterances produced by a same-gender talker. However, perceptually connecting sound across gaps (streaming) can be difficult when sources are too similar, leading to confusions between the competing streams. When spectrotemporal features of simultaneous sounds clearly differ (as in the case of a simultaneous talker and a flute), segregation becomes even easier and streaming happens automatically, driven by continuity of these acoustic features (Bressler et al., 2014).

These automatic, bottom-up processes tend to stream together sounds sharing spectrotemporal properties, even when those features are irrelevant to the listener’s goals. For instance, single-neuron recordings in the brainstem of anesthetized animals show signatures of automatic perceptual organization (Pressnitzer et al., 2008). In the auditory cortex, different neurons respond selectively to different features, like pitch and space, providing a neural substrate for source segregation based on those features (Bizley & Cohen, 2013; Middlebrooks & Waters, 2020). Specifically, some “hard-wired” computations automatically process certain spectrotemporal features, which can lead to distinct neural populations encoding competing sources, providing a foundation for perceptual segregation and selection. Consequently, unattended spectrotemporal features can perceptually “bind” with an attended object that was selected based on some other feature, enhancing streaming and improving recall performance for a target source (Best et al., 2008; Fischer et al., 2024). Thus, spectrotemporal features that may seem irrelevant to a particular task nonetheless automatically contribute to auditory streaming and help listeners process simultaneous sounds.

Building on this foundation, we asked how the continuity of a task-irrelevant feature—voice identity—influences spatial selective attention under interruption. As noted above, continuity of a task-irrelevant feature enhances perceptual segregation, but it may also have other effects. Once a listener has latched onto a target stream based on its location, continuity of a talker’s voice may provide an additional cue to guide attention following an interruption, reducing the cost of interruption (e.g., see Bonacci et al., 2020). Other factors are likely at work as well. Spatial selective attention is not static—it builds gradually over time as the listener continues to focus on the target stream. Distinct talkers may facilitate this buildup by reducing competition between sources, allowing spatial focus to become more stable from syllable to syllable. However, these buildup effects depend on attention remaining properly aligned with the target; when attention drifts or locks onto the wrong stream, the listener must effectively start over on the next syllable (see Bressler et al., 2014). An interruption may automatically “break” this buildup process by resetting spatial focus, instantaneously increasing competition between streams. By resetting buildup, interruptions thus may diminish advantages otherwise conferred by talker differences.

Previous experiments explored spatial selective attention and interruption using a syllable sequence recall task (Liang et al., 2022, 2025). We used the same basic task, presenting two streams of syllables—one target and one distractor—differentiated *only* by spatial location. A random subset of trials contained an interrupter. Listeners were asked to report the syllables from the target stream while ignoring the distractor stream and any interrupter. The experiments documented not only large effects of interruption on recall of the next target syllable, but also persistent errors in recall of even later syllables: refocusing spatial attention after an interruption was difficult when competing streams differed only in location.

Here, we expanded this paradigm by manipulating the similarity of competing stream voices and continuity of voice within streams to test how a task-irrelevant feature shapes attention after interruption. In the Continuous Talker (CT) experiment, each stream presented syllables from a single talker, but the talker could either be the same or different across streams. Experiment CT thus tested whether adding another feature differentiating target and masker reduces the cost of interruption. In the Random Talker (RT) experiment, different talkers could appear within the same stream, disrupting talker continuity. Experiment RT tested how discontinuity of the task-irrelevant talker feature influences performance.

Together, these experiments explored how task-irrelevant acoustic features shape performance during spatial selective attention in the presence of interruptions. We predicted that when target and distractor streams were differentiated by talker identity and the talker was continuous within each sequence, the effect of the interrupter would persist across fewer syllables. Conversely, when talker continuity within a stream was disrupted, we expected weaker stream formation and a bigger interruption cost. Consistent with these predictions, interruption effects were significantly reduced and short-lived when targets and distractors were spoken by different talkers. Continuity of talker identity also produced a clear buildup of benefit, even though attention was always directed to spatial features, demonstrating the role of voice continuity in supporting streaming even when voice identity is an unreliable, task-irrelevant feature.

## II. Methods

### A. Participants

We tested 45 participants (aged 18-72, mean 31.98, std 11.20; 15 females, 28 males, one non-binary, one preferred not to report) for Experiment CT; 44 participants (aged 20-63, mean 35.23, std 9.74; 23 females, 21 males) for Experiment RT. All participants were native English speakers, reported no hearing loss, and had not taken part in previous experiments in this study series. All procedures were approved by the university’s Institutional Review Board. Participants provided informed consent and were compensated for their participation.

### B. Stimuli

In both experiments, participants listened to two competing sound streams simulated as coming from different directions: a target stream and a distractor stream. All sounds were presented through headphones with stereo signals. Sounds were spatialized by convolving mono signals with generic head-related impulse responses (HRIRs) measured from a KEMAR manikin (Gardner & Martin, n.d.). One of the streams was simulated as coming from 30 degrees to the left, the other from 30 degrees to the right.

Each stream consisted of 5 syllables, randomly selected from three consonant-vowel syllables (/ba/, /da/, and /ga/). However, we manipulated the randomization of the target syllables to try to allow us to analyze error patterns. For Experiment CT, the second target syllable and the adjacent, distractor syllables (first and second) were chosen to all differ from one another, so that each of /ba/, /da/, and /ga/ appeared only once across these three syllables. The same was true for target syllable 3 and distractor syllables 2 and 3, and for target syllables 1, 4, and 5. For Experiment RT, target syllables 2, 3, and 4 were chosen to differ from each other so that each of /ba/, /da/, and /ga/ appeared only once in these syllables. In addition, target syllable 1 was chosen to differ from target syllable 2.

We imposed these constraints in an attempt to classify error types, intending to determine whether a mistake reflected a random guess, confusion with the distractor, or confusion with adjacent target syllables. However, in practice, the variety of error patterns was too great to support a meaningful error-type analysis. Nonetheless, because the same constraints were applied consistently within each experiment, comparisons within Experiment CT and within Experiment RT remain valid. In post hoc analyses, we found that the different constraints had systematic effects that complicate direct comparisons across experiments. We describe the consequences of these randomization differences in detail in Section V.

Two separate syllable sets were recorded, one by a male and one by a female native speaker of North American English. Each syllable was windowed to have a duration of 450ms and normalized to have the same root mean square (RMS) level. Syllables were concatenated to form isochronous target and masker streams with a 600ms onset-to-onset interval. The target stream always led and started 300ms before the first distractor syllable, producing temporally interleaved target and distractor streams (Fig. 1).

**Fig. 1.**
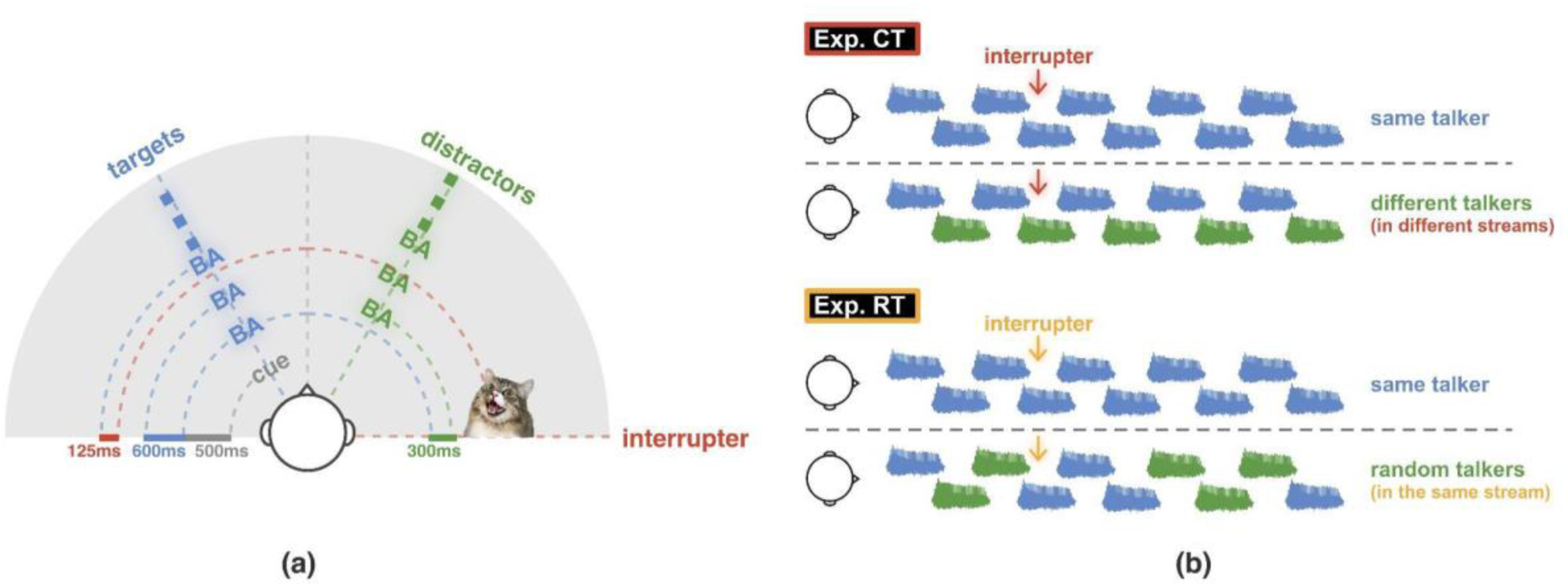
Experiment paradigms. (a) Timing and spatial layout of target and distractor streams and interrupter. (b) Top: Same Talker and Different Talker conditions in Exp CT; bottom: Same Talker and Random Talker conditions in Exp RT.

In Experiment CT, the talker was continuous within each stream, but could either be the same or different across streams. Specifically, both target and distractor streams were spoken by the same talker in 50% of the trials; of these, half presented the male talker and half the female talker (Same Talker condition). In the remaining 50% of trials, the talker differed across streams, with one stream spoken by the male and the other spoken by the female (Different Talker condition). In the Different Talker condition, trials were counterbalanced so that half had the male and half the female as the target, and, independently, half had the male on the left and half on the right.

In experiment RT, half of the trials used the same talker for all syllables (Same Talker condition; half male, half female) as a baseline (identical stimuli to the Same Talker trials in Experiment CT). In the other half of trials, the talker was randomly and independently selected for each individual syllable in both target and distractor streams (Random Talker condition). One spreadsheet was used in Gorilla to control stimuli presentation; thus, the talker randomization was identical across participants.

For both experiments, 50% of the trials in each condition included a 250ms-long interrupting sound 125ms before the onset of the third target syllable. The interrupter was presented from 90 degrees azimuth in the same hemifield as the distractor stream. For each interrupted trial, the interrupting sound was randomly selected from a set of 48 natural sounds (e.g., cat meowing, glass shattering), without replacement. None of the interrupters were human speech and all were perceptually distinct from the syllables. The sounds were retrieved from the internet, windowed to be 250ms in duration, and normalized to have the same RMS level. To ensure their saliency, the level of each interrupter was set 5dB above the syllable level, before spatialization.

Note that the experiments were conducted online, and we were not able to control the overall presentation level. Before the experiments began, the participants were instructed to adjust the sounds to a comfortably loud level.

### C. Task Procedure

Both experiments were conducted online. The online experiments were constructed with the Gorilla platform (gorilla.sc). Data was collected through Prolific (prolific.co).

Headphone screening was performed before the formal experiment to ensure the participants’ playback systems preserved interaural differences in the stereo signals. In the headphone screening task, participants listened to brief broadband noise signals and identified which contained a binaural Huggins pitch (Milne et al., 2021). In each trial, three 1000ms-long noise signals were presented, two of which were diotic and one that imposed a 180 degrees interaural phase shift in a narrow frequency band of otherwise identical noise. This interaural phase manipulation produces a strong pitch perception if and only if the playback system preserved binaural cues. Participants performed six-trial-long blocks of the screening task. Anyone who failed to achieve 100% correct on a block within three tries was rejected.

Participants were instructed to maintain their gaze on a fixation cross at the center of the screen during the stimulus presentation. At the beginning of each trial, an auditory cue (/ba/) from either 30 degrees to the left or to the right cued the direction of the target on that trial. The target stream started 500ms after the offset of the cue. Participants were instructed to attend to the syllable stream coming from the target direction while ignoring the competing distractor stream and any interrupting sounds. After the stimulus presentation, the participants were asked to click on a graphical user interface on their computer screen to report the five syllables (in order) that they heard from the target direction.

Prior to formal data collection, participants participated in 6-trial-long training blocks. The training trials were identical to uninterrupted trials in the experiments and were spoken by the same male talker. Feedback was provided after each trial throughout the training session, indicating the correct target sequence. Participants who failed to report 4 out of 5 syllables correctly in at least 3 out of 6 trials within five tries were rejected.

Participants proceeded to the task session after successfully completing the headphone screening and training session. There were 96 trials in each task session, divided into two blocks of 48 trials. Trials were randomized within blocks. In each block, half of the trials were uninterrupted and half interrupted. Of the 24 uninterrupted trials in a block, half were the Same Talker condition and the half the other talker condition (Different Talker in experiment CT; Random Talker in experiment RT). Similarly, half of the interrupted trials were Same Talker and half the other talker condition.

### D. Data analysis

For both experiments, each participants’ percent correct syllable recall performance was computed by averaging scores across trials within each condition, separately for each syllable position. The interruption effect was then computed as the percent correct syllable recall in uninterrupted trials minus that in interrupted trials, computed separately for each participant and then averaged across participants. Similarly, in Experiment CT, we computed the different talker benefit as the percent correct syllable recall for the Different Talker condition minus that for the Same Talker condition; in Experiment RT, we computed the continuous talker benefit as the percent correct syllable recall for the Same Talker condition minus that for the Random Talker condition.

For both experiments, statistical tests were conducted only on the interruption effect and the different talker and continuous talker effects. Although we show plots of the raw percent correct syllable recall, we do not conduct statistical tests on the raw data to avoid redundancy. Holm-Bonferroni corrected t-tests were first performed on both the interruption effect and the two talker effects to examine whether these effects were greater than zero at each distinct syllable position. We then undertook repeated-measure ANOVA with main factors of talker condition (same vs. different for Experiment CT; same vs. random for Experiment RT) and syllable position (1-5) on interruption effect data. For significant main effects and interaction effects, post hoc Holm-Bonferroni corrected pairwise comparisons were conducted. Similarly, repeated-measure ANOVA and post hoc comparisons were performed for talker effect, with main factors of interruption condition (uninterrupted, interrupted) and syllable position (1-5).

For these sequence recall tasks, we also analyzed the effect of past history to determine whether recall performance for a syllable differed depending on whether or not participants correctly reported the previous syllable. Specifically, we looked to see if the likelihood of correctly reporting a syllable was greater when a listener properly reported the previous syllable compared to when they got the previous syllable wrong. Overall, such dependencies are likely to matter: if a participant’s attention strays during a trial, the effect is likely to last beyond one syllable. The question we were most interested in was whether these conditional probabilities depended on the talker condition (same or different in Experiment CT, same or random in Experiment RT).

For both experiments, we first computed the raw percent correct performance for each syllable position, broken down by whether the previous target syllable was reported correctly or incorrectly. We then computed the “benefit of previous syllable correct” as the percent correct performance when the previous syllable was reported correctly minus the percent correct performance when the previous syllable was incorrectly recalled. We then performed Holm-Bonferroni corrected t-tests to determine whether this difference was significantly greater than zero for syllables 2-5. We also fit a linear mixed-effect model with fixed effects of talker conditions (same vs. different for Experiment CT; same vs. random for Experiment RT) and syllable positions (2-5) and random effect of subject with random intercepts for each interruption condition. Post hoc Tukey-adjusted pairwise comparisons were conducted when significant effects were found.

For Experiment RT, in the Random Talker condition, we further broke down recall performance by whether or not the previous target syllable was spoken by the same talker as the current target syllable. We contrasted performance for these two cases and compared them to the Same Talker condition. We then fit a linear mixed-effect model with fixed effects of talker conditions (Same Talker, Random Talker - Same, Random Talker - Different) and syllable positions (2-5) and a random effect of subject with random intercepts for each interruption condition. Post hoc Tukey-adjusted pairwise comparisons were conducted to help interpret all significant effects.

## III. Continuous Talker (CT)

### A. Results

This experiment compares syllable recall performance and interruption effects with the same or different talkers for target and distractor streams. Figure 2a shows the raw percent correct syllable recall rate for each syllable position in the sequences for all four task conditions. In most of the syllable positions, listeners perform better in recalling the targets when the target and distractor streams are spoken by different talkers (unfilled bars) than the same talker (filled bars). The interrupter, which occurred just before the third target syllable in half of the trials, resulted in lower syllable recall performance for interrupted trials (red) than uninterrupted trials (blue) for the third target syllable. For the subsequent 4th and 5th target syllables, the interrupter reduced recall performance when the target and masker streams were the same talker (filled bars), replicating earlier reports (Liang et al., 2022, 2025); however, these later syllables showed little impact of the interrupter when the streams were spoken by different talkers (unfilled red and blue bars are similar for the 4th and 5th syllables).

**Fig. 2.**
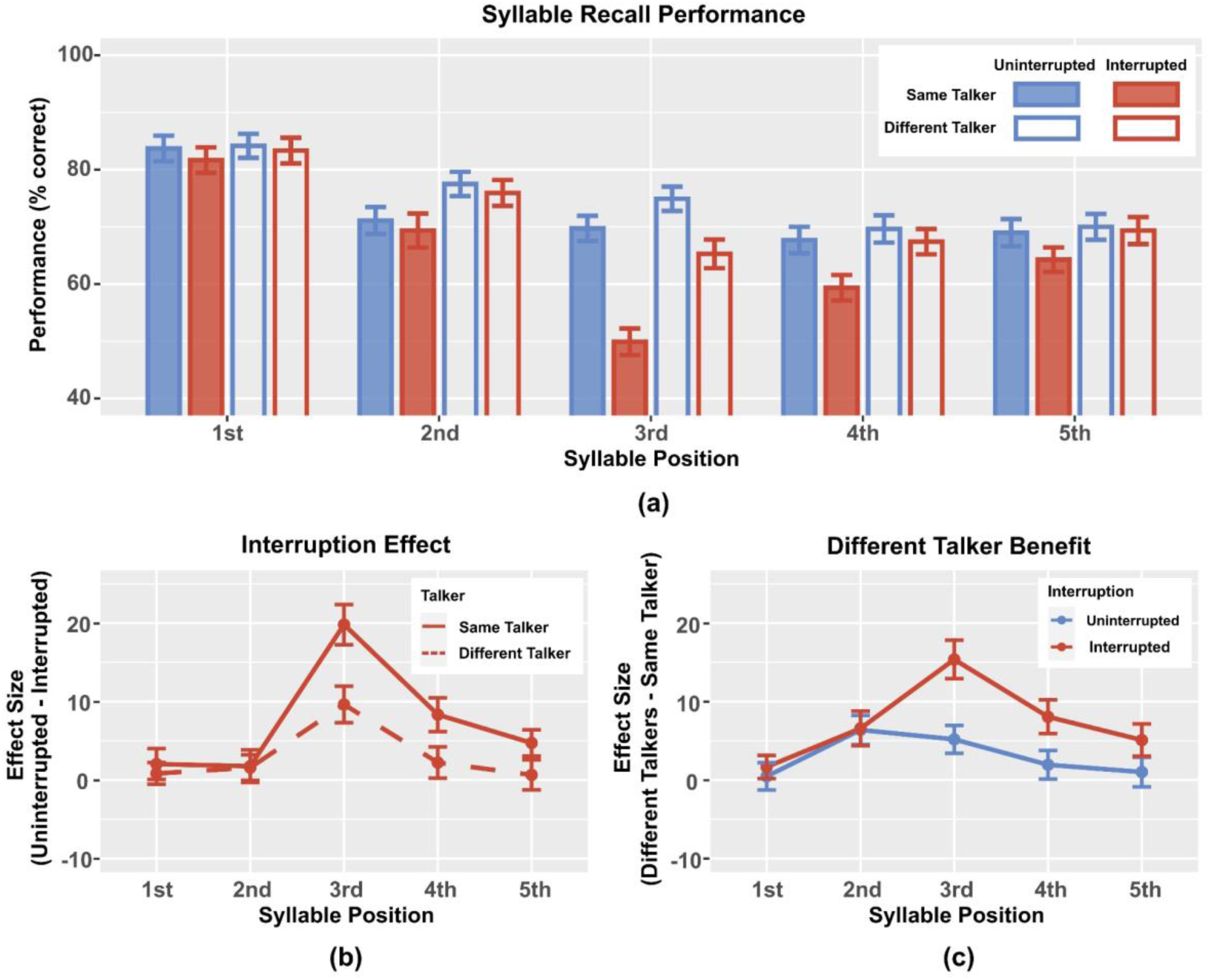
(a) Mean syllable recall performance, averaged across participants. Blue represents uninterrupted trials and red interrupted trials; filled bars represent Same Talker conditions and unfilled bars Different Talker conditions. Error bars show the across-participant standard error of the mean. (b) Interruption effect for the Same Talker and Different Talker conditions, computed as percent correct performance in the uninterrupted trials minus that in the interrupted trials, averaged across participants. Error bars show the across-participant standard error of the mean. (c) Talker effects in uninterrupted and interrupted trials, computed as percent correct performance in the Different Talker condition minus that in the Same Talker condition, averaged across participants. Error bars show the across-participant standard error of the mean.

#### 1. Interruption effect

To highlight the impact of interrupter on syllable recall, Figure 2b shows the interruption effect, computed as the percent correct syllable recall performance in uninterrupted trials minus that in interrupted trials, plotted as a function of syllable position. The interrupter occurred before the third target, which shows the largest interruption effect (around 20% for the Same Talker condition and 10% for the Different Talker condition). For the Same Talker condition, the interruption effect was smaller but positive for syllables 4 (∼8%) and 5 (∼5%), but for the Different Talker condition, the interruption effect was much smaller.

Statistical analyses confirmed these observations. The interruption effect was not significantly greater than zero for either same or different talker conditions for syllables 1 and 2. However, the interruption effect for syllable 3 was significant for both same and different talker conditions(*t*_44_ = 7.74, *p* < 0.001 and *t*_44_ = 4.15, *p* < 0.001, respectively). For the Same Talker condition, the interruption effect was significant for both target syllable 4 (*t*_44_ = 3.87, *p* = 0.001) and syllable 5 (*t*_44_ = 2.81, *p* = 0.026), but for the Different Talker condition, there was no statistically significant impact of the interrupter on these syllables.

We ran a repeated-measure ANOVA on the interruption effect with main factors of talker condition (same, different) and syllable position (1-5). There was a significant effect of talker condition, confirming that the interruption effect was larger in the Same Talker condition than the Different Talker condition (*F*_1,44_ = 6.962, *p* = 0.011). Syllable position was also significant (*F*_4,176_ = 17.894, *p* < 0.001), as was the interaction between talker condition and syllable position (*F*_4,176_ = 3.154 *p* = 0.016). Post hoc tests showed that the interruption effect was larger for the Same Talker than Different Talker condition for syllable 3 (*t*_44_ = 3.648, *p* = 0.003). In the Same Talker condition, the interruption effect on syllable 3 was significantly larger than for all other syllable positions (syllable 1: *t*_44_ = 5.465, *p* < 0.001; syllable 2: *t*_44_ = 6.178, *p* < 0.001; syllable 4: *t*_44_ = 4.166, *p* = 0.022; syllable 5: *t*_44_ = 5.195, *p* < 0.001). In contrast, after multiple-comparison correction, the interruption effect did not differ significantly between any of the pairs of syllable positions in the Different Talker condition.

#### 2. Talker effect

Figure 2c shows the talker effect, computed as percent correct performance in the Different Talker condition minus that in the Same Talker condition, quantifying the benefit of having distractors spoken by a different talker. In general, the talker effect was positive, demonstrating a performance advantage when the streams were spoken by distinct talkers. In the uninterrupted condition, the talker benefit was modest, ranging from near zero for the first and fifth syllable to about 5% for the second and third syllables. For syllables 3-5, the benefit of having different talkers was larger when the interrupter was present than in uninterrupted trials. The talker effect was greatest, and the difference between the talker effect in the interrupted and uninterrupted conditions was greatest, for syllable 3, which occurred right after the interrupter.

T-tests showed that the talker effect was significantly greater than zero for all but the first syllable in the interrupted condition (syllable 2: *t*_44_ = 2.99, *p* = 0.016; syllable 3: *t*_44_ = 6.24, *p* < 0.001; syllable 4: *t*_44_ = 3.73, *p* = 0.002; syllable 5: *t*_44_ = 2.47, *p* = 0.044). In the uninterrupted condition, the talker effect was significant only for syllables 2 and 3 (*t*_44_ = 3.43, *p* = 0.005 and *t*_44_ = 2.93, *p* = 0.016, respectively).

Repeated-measure ANOVA on the talker effect with factors of interruption condition (uninterrupted, interrupted) and syllable position (1-5) found that both main effects were significant (*F*_1,44_ = 6.962, *p* = 0.011 and *F*_4,176_ = 9.135, *p* < 0.001, respectively); their interaction was also significant (*F*_4,176_ = 3.154, *p* = 0.016). Post hoc tests showed that on the third target syllable, the talker effect was larger for interrupted than uninterrupted trials (*t*_44_ = 3.648, *p* < 0.001), but the talker effect was not significantly different between interrupted and uninterrupted trials for any other syllable positions. In the interrupted condition, performance for the third syllable showed a significantly larger talker effect than on the first syllable (*t*_44_ = 5.290, *p* < 0.001); however, the talker effect was not significantly different between any other syllable pairs in the interrupted condition or any syllable pairs in the uninterrupted condition.

#### 3. Within-trial effect of performance on previous syllable

For both uninterrupted and interrupted trials, and across all syllable positions, there was a strong sequential effect: participants were more likely to correctly recall a target when the previous syllable was correctly recalled than when it was incorrectly recalled (Fig. 3 top panels: solid bars are taller than corresponding striped bars for all pairs).

**Fig. 3.**
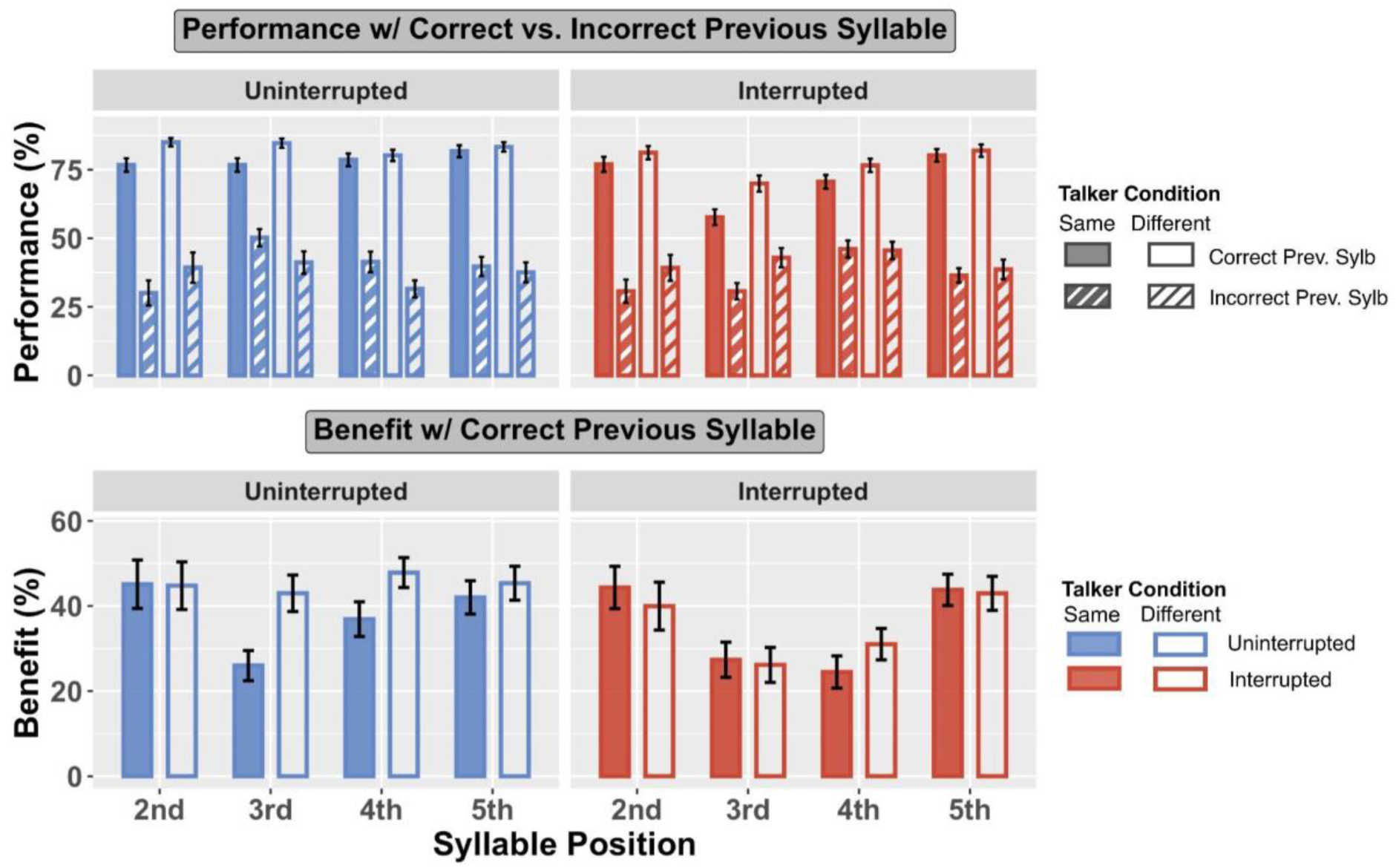
Top panels: Performance with previous syllable correctly vs. incorrectly recalled. Bottom panels: Benefit of having previous syllable correctly recalled, computed as the performance with previous syllable correctly recalled minus performance when those were incorrectly recalled.

There is another consistent effect for both uninterrupted and interrupted trials: when the previous syllable was heard correctly (solid bars), performance was always better for the Different Talker condition than for the Same Talker condition (the open solid bars are always higher than the corresponding filled solid bars in both panels). However, when listeners get the previous syllable incorrect (striped bars), performance differs for uninterrupted and interrupted trials. Specifically, when the previous syllable is incorrect in the uninterrupted trials, performance in the Different Talker condition tends to be slightly worse than for the Same Talker (in the left panel of Fig. 3, the white bars with thin blue stripes are generally lower than the blue bars with thin white stripes); however, for interrupted trials, performance in the Different Talker condition tends to be slightly better than for the Same Talker condition (in the right panel of Fig. 3, the white bars with thin reddish stripes are generally higher than or equal to the reddish stripes with thin white stripes).

To quantify whether the benefit of correctly reporting a previous syllable varies with talker condition and interruption, we computed the average “benefit of previous syllable correct” (percent correct on a syllable when the previous syllable was correctly recalled minus that when the previous syllable was incorrectly recalled). The bottom panels of Figure 3 show this benefit separately for uninterrupted and interrupted trials (left and right panels, respectively), and for same talker and different talker (filled and open bars, respectively).

In uninterrupted trials, the benefit of getting the previous syllable correct tends to be greater in the Different Talker condition than the Same Talker condition (in the bottom left panel of Fig. 3, open bars are taller than filled bars for target syllables 3, 4, and 5). However, in interrupted trials, no such pattern appears; instead, the benefit of getting the previous syllable correct varies with syllable position. Specifically, the benefit is small for target syllable 3, which was right after interrupter, and the subsequent syllable (target syllable 4).

T-tests confirmed that the benefit of getting the previous syllable correct was positive for all syllable positions (*p* < 0.001 for all). Linear mixed effect models with fixed effects of talker conditions (same, different) and syllable positions (1-5) and a random effect of subject were fit for the previous-benefit data, separately for uninterrupted and interrupted conditions.

For uninterrupted trials, there was a significant greater previous syllable correct benefit in the Different Talker than the Same Talker condition (*F*_1,311.01_ = 6.511, *p* = 0.011), but no significant effect for syllable positions and no significant interaction between talker condition and syllable position. Thus, in the uninterrupted trials, the benefit of properly attending and reporting the previous syllable had a larger effect on performance for Different Talker than for Same Talker trials.

For interrupted trials, there was a significant effect of syllable position (*F*_3,314.65_ = 12.404, *p* < 0.001), but not for talker condition and not for the interaction of syllable position and talker condition. Post hoc pairwise comparisons showed the benefit on target syllable 2 was significantly greater than that on target syllable 3 (*p* < 0.001) and target syllable 4 (*p* = 0.001); also, the benefit on target syllable 5 was significantly greater than that on target syllable 3 and target syllable 4 (*p* < 0.001 for both). There were not significant differences between the benefits for syllables 2 and 5 or for syllables 3 and 4. Thus, for interrupted trials, the benefit of properly attending and reporting the previous syllable was smaller for the syllable immediately after and two after the interruption than for the other syllables.

### B. Discussion

Experiment CT investigated how differences in talker identity between target and distractor streams influence spatial selective attention when a salient, task-irrelevant sound interrupts attention. Specifically, we asked whether presenting competing streams with different talkers reduced the disruptive impact of an interrupter and improved recall of target syllables, even though spatial location alone defined the target stream.

Three key findings emerged. First, when the target and distractor were spoken by different talkers, listeners recalled target syllables more accurately overall. Second, the interrupter impaired recall of the third target syllable, which occurred 150ms after the interrupter, in all conditions, consistent with the interrupter involuntarily capturing attention; however, this decrement was significantly smaller in the Different Talker condition than in the Same Talker condition. Third, the interrupter produced a long-lasting decrement in the Same Talker condition, persisting through the fourth and fifth target syllables. However, when the two streams differed by voice, only recall of the third target syllable was affected by the interrupter.

The observed pattern of results suggests that talker differences benefit performance through three complementary mechanisms. First, talker differences enhance perceptual segregation and streaming. Second, talker identity differentiates target and masker syllables stored in working memory. Third, talker-based attention augments or even supplants spatial attention following an interruption.

We also observed sequential effects: listeners were more likely to report a syllable correctly when they heard the previous syllable correctly. Importantly, this benefit of correctly recalling the previous syllable showed different patterns in uninterrupted versus interrupted trials. In uninterrupted trials, the benefit was greater for Same Talker condition than for Different Talker condition; but in interrupted trials, the benefit did not differ with talker condition. Instead, in interrupted trials, the benefit varied with syllable position—it was small for syllables after the interruption. These differences show that the benefit of getting the previous syllable correct does not simply reflect trial-to-trial fluctuations in attentiveness, but also depends on stimulus characteristics (uninterrupted vs. interrupted, Same Talker vs. Different Talker), reflecting effects of perceptual continuity of the auditory stream on performance.

#### 1. Talker differences support perceptual segregation and streaming

Even in uninterrupted conditions, talker differences provided small but significant improvements in target recall compared to when the talkers were the same. Although talker identity covaries with spatial location and thus may appear redundant, talker differences nonetheless strengthen perceptual streaming and improve selective attention. In addition, talker differences supply an additional cue for maintaining attentional selection; past work shows that even when a target stream is defined by its location, listeners may use more robust spectrotemporal features to sustain attention to an ongoing target stream (Bonacci et al., 2020). Because male and female voices differ systematically in pitch and spectral composition, their neural representations overlap less in the auditory cortex, making it easier to maintain separate object representations for target and distractor streams. Such improvements can account for the small but consistent advantage observed in the Different Talker condition during uninterrupted trials.

#### 2. Talker cues differentiate syllables recalled from working memory

The large drop in accuracy for recall of the third target syllable likely reflects a transient disruption of attentional focus—a hijacking of top-down attention by the salient interrupt away from the target stream. When attention is diverted by the interrupter, the third target syllable must often be reconstructed from working memory rather than perceived directly. In working memory, auditory object representations bind together spectrotemporal features that not only define syllable content, but also include pitch, timbre, and other voice-related features (Bizley & Cohen, 2013). Our results suggest that spatial location is only weakly and inconsistently bound to spectrotemporal features of an auditory object in working memory, explaining why the cost of interruption is much larger in the Same Talker compared to Different Talker conditions.

Past work helps explain why this could be the case. Spatial attributes of auditory events are represented only weakly in the auditory cortex unless listeners are actively engaged in a task requiring spatial information (Lee & Middlebrooks, 2011). Such auditory spatial tasks strongly recruit a frontoparietal visuospatial cognitive control network that is not inherently auditory (Michalka et al., 2015). Disruptions of top-down spatial attention likely interfere with volitional engagement of the visuospatial network necessary for spatial auditory processing, so that working memory fails to bind spatial to spectrotemporal attributes. Consequently, when target and distractor share the same talker, spatial cues alone cannot robustly guide retrieval of the correct syllable from working memory, increasing confusions and recall errors. In contrast, when voices differ, talker identity acts as an intrinsic retrieval cue, enabling listeners to sort stored syllables by voice and recover the target more accurately.

#### 3. Talker identity provides an alternative feature to guide top-down attention

The effects of the interrupter persisted through the fourth and fifth syllables only in the Same Talker condition. When the target and distractor voices differed, performance on these syllables was statistically indistinguishable from uninterrupted performance, suggesting that listeners could re-establish focus on the target stream more efficiently. This pattern indicates that talker cues not only facilitate streaming but also aid the reorientation of top-down attention following an interrupter. When target and distractor talkers differ, after attention is drawn away, listeners can refocus not only on where the target is located but also on who is speaking—using talker identity as a stable feature to guide attention.

This explanation echoes prior results showing that in spatial selective-attention tasks (where location defines the target), listeners may shift to maintaining attention based on other object-defining features such as pitch, when available (Bonacci et al., 2020). In the absence of such cues, as in the Same Talker condition, refocusing relies solely on spatial information, which appears to be slower and more error-prone. The persistence of the interruption effect in the Same Talker condition through the fourth and fifth syllables likely reflects the gradual buildup of selective attention when a listener maintains focus on the same spatial location over time. For instance, when listeners sustain top-down spatial attention to a stable target location, performance in a spatial selective attention task improves from syllable to syllable (Best et al., 2008). This kind of slow buildup may reflect the need for auditory spatial attention to recruit the non-native visuospatial network that supports spatial target selection. In the Different Talker condition, attention to talker identity, established in the first and second syllables, can bypass this slower spatial-control system, reorienting attention rapidly following an interrupter.

#### 4. Interruptions break perceptual effects of stream continuity

Sequential dependencies in performance across syllables revealed another influence of talker continuity. In general, listeners were more accurate when they had correctly identified a preceding syllable than when they had responded incorrectly on the previous syllable. In uninterrupted trials, this sequential dependency differed between talker conditions, reflecting differences in whether the two competing streams were perceptually distinct or spectrotemporally similar. However, when an interruption occurred, any influence of stream continuity disappeared; instead, performance depended only on the temporal proximity of the syllable to the interruption itself.

In uninterrupted trials, the benefit of having correctly identified the previous syllable was larger in the Different Talker condition, where the two competing streams were spectrotemporally distinct, than in the Same Talker condition, where they differed only in location. Past studies show that when listeners focus attention on one auditory element, a subsequent element that is spectrotemporally similar is automatically more likely to capture attention (Bressler et al., 2014). This aligns with the current results: when listeners were correct on the previous syllable, automatic streaming, not spatial attention alone, increased the likelihood of maintaining attention on the same spectrotemporal stream in Different Talker trials. Conversely, when listeners misidentified a syllable—often by incorrectly attending to the distractor stream—that same mechanism may have reinforced attention to the wrong voice, reducing accuracy on the next syllable. Together, results from the uninterrupted trials suggest that continuity of distinct target and masker streams contributes strongly to listeners’ ability to maintain focus and resist interference.

A salient interruption, however, broke perceptual continuity. The interruption effectively “reset” the buildup of streaming, causing the benefit of getting the previous syllable correct to nearly vanish for subsequent syllables. Only by the final syllable did the prior-correct benefit begin to re-emerge. Thus, while perceptual and attentional buildup strengthen over time, they remain vulnerable to disruption by salient, attention-capturing events.

#### 5. Conclusions from Experiment CT

Taken together, these findings replicate prior work showing that unattended features bind with an attended object, improving target selection and recall in selective attention tasks (e.g., Best et al., 2008; Fischer et al., 2024). They extend that work by demonstrating that voice differences between concurrent streams mitigate the disruptive effects of a salient interrupter during spatial selective attention. These results demonstrate that spatial selective attention in complex auditory scenes is supported by a dynamic interplay among perceptual, mnemonic, and attentional-control mechanisms—all of which can be reinforced by consistent non-spatial cues such as voice identity. Sequential analyses show that the continuity of distinct voices reinforces perceptual streaming, allowing attentional focus to build up over time. However, this buildup is fragile, easily broken by salient events that disrupt top-down attention. Thus, distinct voice cues can enhance resistance to disruption and allow streaming to build over time but cannot fully overcome the involuntary capture of attention by new events.

## IV. Random Talker (RT)

### A. Results

Experiment RT compares performance when both target and distractor streams were spoken by the same talker with performance when the talker for each syllable within each stream was randomized, independently, within a trial. We expected talker randomization to break the continuity of acoustic features for both streams and make the task harder. Consistent with this prediction, performance in the Random Talker condition generally led to lower syllable recall performance than Same Talker (Fig. 4a, unfilled blue and yellow bars are generally lower than the corresponding filled ones). As in many of our previous experiments in this series, performance on the first and last syllables was typically better than that for the middle syllables, reflecting primacy and recency effects in recall.

**Fig. 4.**
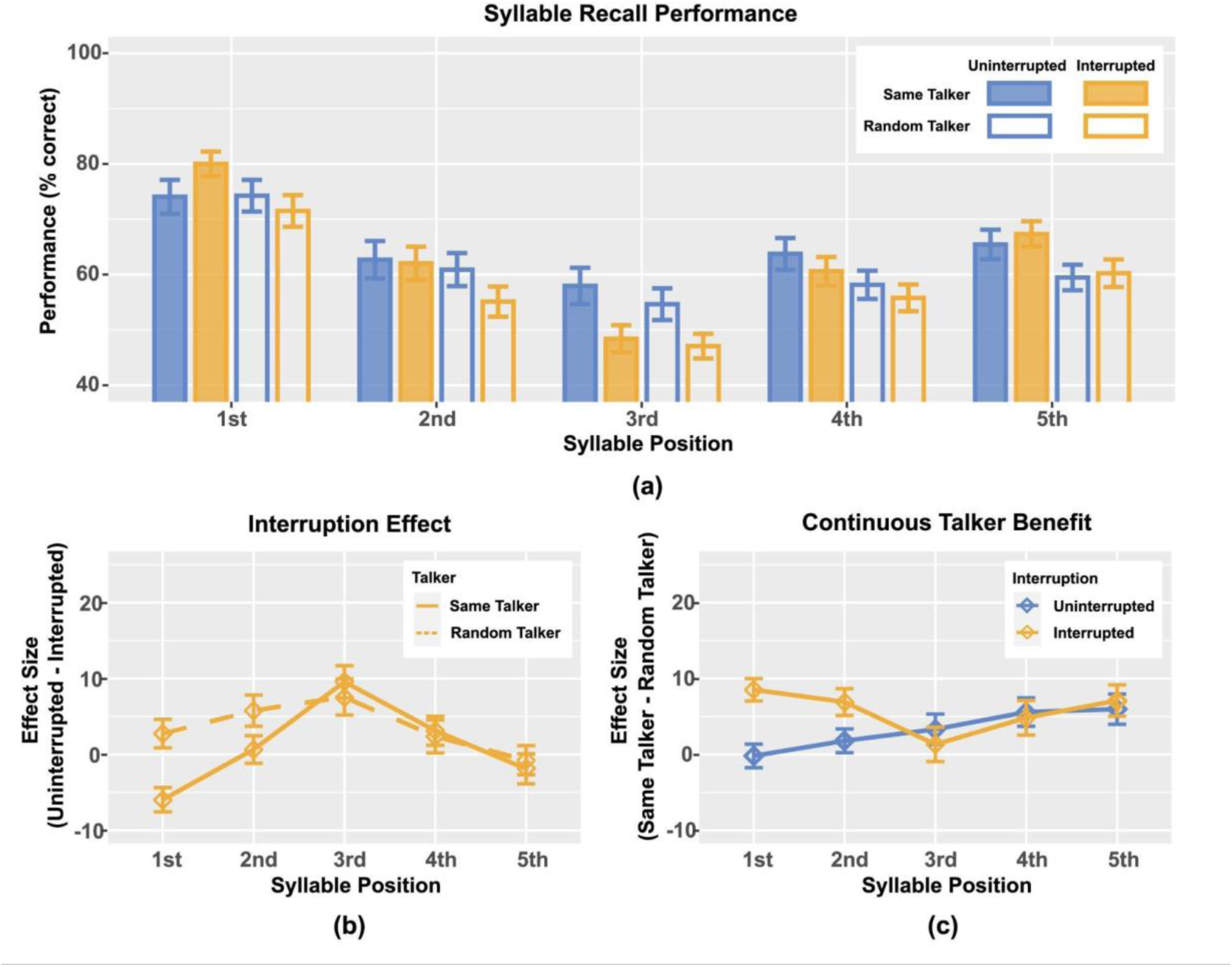
(a) Mean syllable recall performance in Experiment RT, averaged across participants. Blue represents uninterrupted trials and yellow interrupted trials; filled bars show same talker conditions and unfilled bars random talker conditions. (b) Interruption effects for the same and random talker conditions, computed as percent correct in the uninterrupted trials minus that in the interrupted trials, averaged across participants. (c) Talker effects in uninterrupted and interrupted trials, computed as percent correct performance in the same talker condition minus that in the random talker conditions, averaged across participants. Error bars show the across-participant standard error.

#### 1. Interruption effect

We computed the interruption effect (Fig. 4b) as the average percent correct performance in the uninterrupted trials (blue values in Fig. 4a) minus that in the corresponding interrupted trials (yellow values in Fig. 4a). The interrupter always occurred before the third target syllable, leading to a large interruption effect for both Same and Random Talker conditions for syllable 3. The interruption effect was similar for Same and Random Talker conditions for both the fourth syllable, where there was a modest effect, and the fifth syllable, where the effect was near zero. In the Random Talker condition, the average interruption effect was greater than zero for both the first and second syllables. Unexpectedly and unlike all previous results in this series of studies, in the Same Talker condition, listeners performed *better* in interrupted trials than uninterrupted trials for the very first target syllable, producing a *negative* interruption effect for this syllable; the interruption effect was near zero for the second syllable in the Same Talker condition.

These observations were supported by statistical analyses. Two-sided t-tests showed that there was a significant positive interruption effect for the Random Talker on target syllable 2 (*t*_43_ = 2.81, *p* = 0.052) and target syllable 3 (*t*_43_ = 3.20, *p* = 0.021); and for the Same Talker on target syllable 3 (*t*_43_ = 4.50, *p* < 0.001). There was also a significant *negative* interruption effect on target syllable 1 (*t*_43_ = −3.76, *p* = 0.005) with the Same Talker condition. The interruption effect was not significantly different from zero for any other syllable in either Same or Random Talker conditions.

We have never previously seen performance to be better for *any* syllable for interrupted trials compared to uninterrupted trials. We therefore examined performance on syllable 1 to understand whether this was a statistical fluke. Of the 44 subjects, 25 performed better on syllable 1 for interrupted trials than for uninterrupted trials (12 performed better in uninterrupted trials, 7 equally performed in uninterrupted and interrupted), confirming that the effect is consistent (see Supplemental Figure S1). This result is considered further in Section V, which considers differences between Experiments CT and RT.

Repeated-measure ANOVA with main factors talker (same, random) and target syllable (1-5) showed a significant main effect of syllable position (*F*_4,172_ = 8.427, *p* < 0.001) but not talker condition (*F*_1,43_ = 3.471, *p* = 0.069), and a significant interaction between syllable position and talker condition (*F*_4,172_ = 3.853, *p* = 0.007). Post hoc pairwise comparisons showed a significantly greater interruption effect on the first target syllable with Random Talker than Same Talker (*t*_43_ = 4.144, *p* < 0.001), and no significant difference between those two talker conditions at other syllable positions. In the Same Talker condition, the interruption effect on the third target syllable was significantly larger than that on the first (*t*_43_ = 7.57, *p* < 0.001), the second (*t*_43_ = 4.02, *p* = 0.034), and the fifth target syllable (*t*_43_ = 4.09, *p* = 0.031). The interruption effect on the fourth target was also significantly greater than that on the first target syllable (*t*_43_ = 4.06, *p* = 0.033).

#### 2. Talker effect

The benefit of keeping the talker the same rather than randomized (computed as performance in the Same Talker condition minus that in the Random Talker condition) differed between uninterrupted and interrupted trials (see Fig. 4c). In the uninterrupted trials (blue in Fig. 4c), the benefit of the talker being continuous increased monotonically with syllable position. In the interrupted trials, the benefit was similar to that in the uninterrupted trials for syllables 3, 4, and 5; however, there was a much greater talker benefit for the two syllables before the interrupter, with subjects performing roughly 8% better in the Same Talker condition than when the talker was randomized.

Single-sided t-tests showed that, in uninterrupted trials, the talker benefit was greater than zero for target syllable 4 (*t*_43_ = 2.99, *p* = 0.016) and target syllable 5 (*t*_43_ = 2.98, *p* = 0.016). In interrupted trials, the talker benefit was greater than zero for target syllable 1 (*t*_43_ = 5.79, *p* < 0.001), target syllable 2 (*t*_43_ = 3.94, *p* = 0.001), and target syllable 5 (*t*_43_ = 3.45, *p* = 0.005).

Repeated-measure ANOVA with main factors of interruption condition (uninterrupted, interrupted) and syllable position (1-5) showed no significant main effects, but a significant interaction effect (*F*_4,172_ = 3.853, *p* = 0.005). Post hoc pairwise comparisons showed that interrupted trials had a significantly greater talker benefit than uninterrupted trials on target syllable 1 (*t*_43_ = 4.14, *p* < 0.001). The talker benefit was not significantly different across syllable positions within each interruption condition.

#### 3. Within-trial effect of performance on previous syllable

Similar to Experiment CT, participants were more likely to correctly recall a target when the previous syllable was correctly recalled than when it was incorrectly recalled (Fig. 5 top panels: solid bars are taller than corresponding striped bars for all conditions). For both uninterrupted and interrupted trials (left and right panels of Fig. 5, respectively), performance was consistently better for the Same Talker condition than Random Talker condition when the previous syllable was correctly recalled (solid filled bars are always taller than the corresponding open bars). However, there was no consistent effect when the previous syllable was incorrect (the two striped bars are similar in height at each syllable position).

**Fig. 5.**
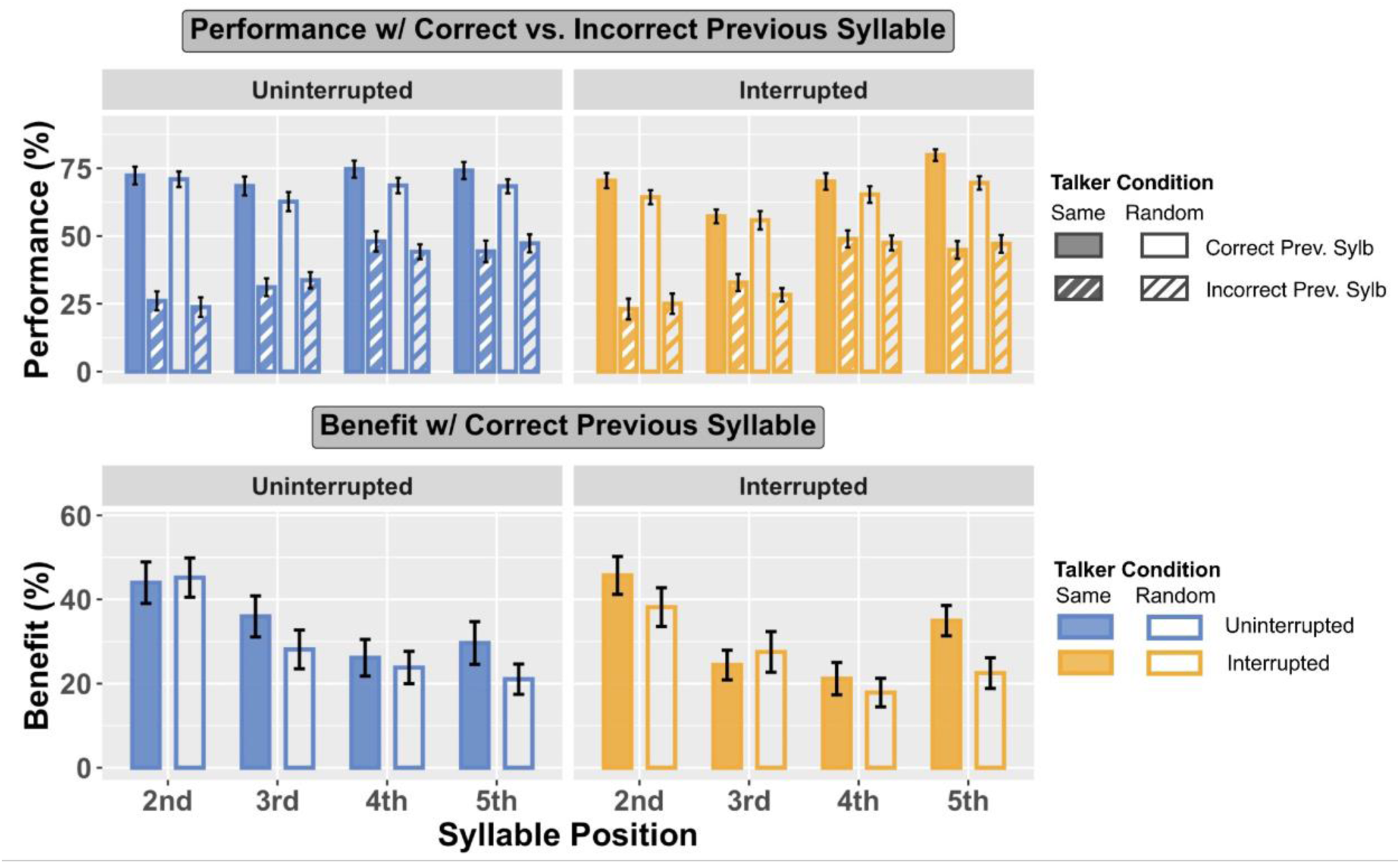
Top panels: Performance with previous syllable correctly vs. incorrectly recalled. Bottom panels: Benefit of having the previous syllable correctly recalled, computed as the performance with previous syllable correctly recalled minus performance when those were incorrectly recalled.

Syllable position had a greater effect on performance when the previous syllable was incorrectly reported than when it was correctly reported. For both uninterrupted and interrupted trials, performance when the previous syllable was incorrect increased from about 20% correct for syllable 2 to about 50% correct for syllable 5. When the previous syllable was correctly reported, performance varied little with syllable position in uninterrupted trials, while for interrupted trials, performance was lowest for syllable 3 (after the interruption) and highest for syllable 5.

The benefit of correctly reporting the previous syllable, computed as the percent correct syllable recall on a syllable when the previous syllable was correctly recalled minus that when the previous syllable was incorrectly recalled, is plotted in the bottom panels of Figure 5. For both uninterrupted (bottom left panel) and interrupted (bottom right panel) conditions, the benefit of getting the previous syllable correct was greatest for the second syllable position. For uninterrupted trials, this benefit generally decreased with syllable position. For interrupted trials, the benefit was relatively small for syllables 3 and 4 and larger for syllables 2 and 5, especially in the Same Talker trials (filled bars).

T-tests confirmed that the benefit of getting the previous syllable correct was positive for all syllable positions in all conditions (*p* < 0.001 for all). Linear mixed effect models with fixed effects of talker conditions (same, random) and syllable positions (2-5) and a random effect of subject were fit for the previous syllable correct benefit data, separately for uninterrupted and interrupted conditions.

For uninterrupted trials (bottom left panel of Figure 5), there was a significant effect of syllable position (*F*_3,287.48_ = 9.566, *p* < 0.001), but no significant effect of talker or interaction between syllable position and talker condition. Post hoc pairwise comparisons showed that the benefit on target syllable 2 was significantly greater than that on target syllable 3 (*p* = 0.012), target syllable 4 (*p* < 0.001), and target syllable 5 (*p* < 0.001); there were no significant differences between target syllables 3, 4, and 5.

For interrupted trials (bottom right panel of Figure 5), there were significant main effects of syllable position (*F*_3,295.07_ = 13.423, *p* < 0.001) and small effect of talker condition (*F*_1,294.33_ = 3.953, *p* = 0.048), but no significant interaction between them. Post hoc pairwise comparisons showed the benefit on target syllable 2 was significantly greater than that on target syllable 3 (*p* < 0.001), target syllable 4 (*p* < 0.001), and target syllable 5 (*p* = 0.002); the benefit on target syllable 5 was significantly greater than that on target syllable 4 (*p* = 0.048), with no other significant differences between target syllable positions.

#### 4. Within-trial effect of talker continuity

In Random Talker trials, the talker was selected independently for each syllable. If the effects of talker continuity are automatic, this should impact performance. To explore this possibility, we analyzed performance based on whether a target syllable was spoken by the same talker as the previous target syllable (Fig. 6), both for Uninterrupted trials (left panel) and for Interrupted trials (right panel). Within each panel, we compare trials from the Same Talker condition (filled bars), where the talker was the same for all syllables (both target and distractor), to trials from the Random Talker condition in which the previous target syllable was the same talker as the current syllable (henceforth, Random Talker - Same condition; open bars) and Random Talker trials where the previous syllable and current syllable were spoken by different talkers (Random Talker - Different condition; striped bars).

**Fig. 6.**
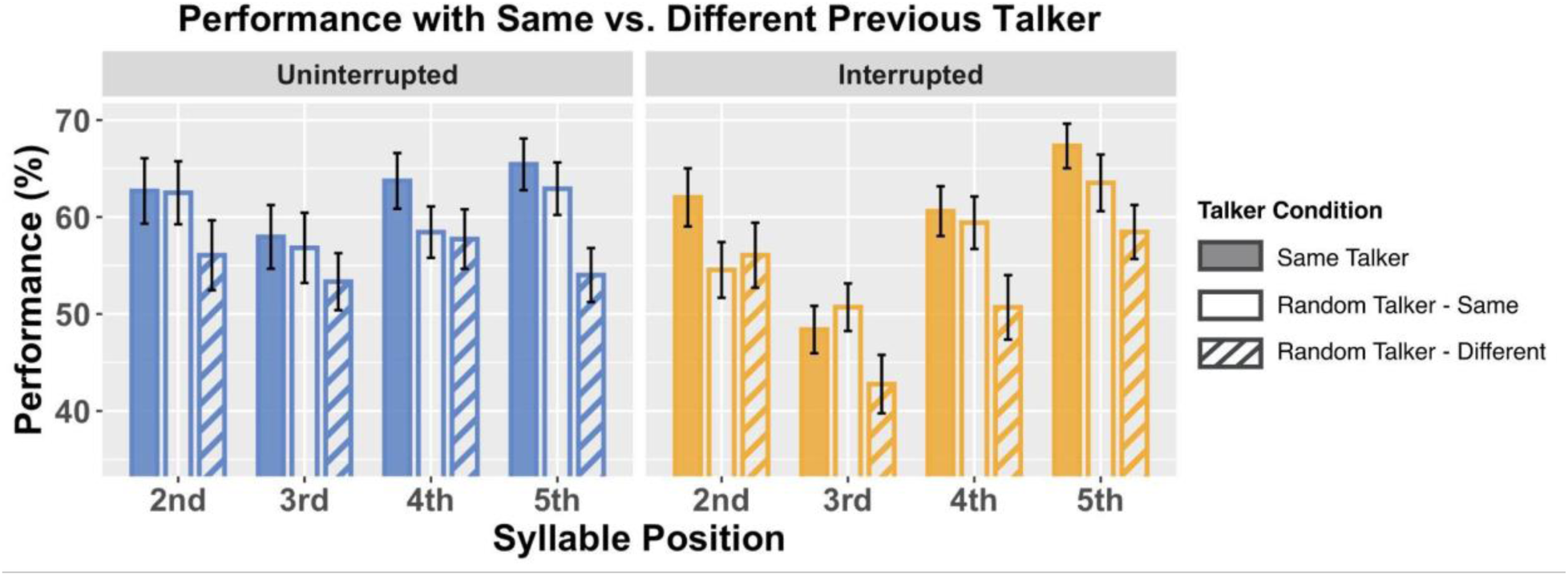
Performance grouped by talker of the previous syllable, with Same Talker vs. Random Talker - Same (talker unchanged from previous target) vs. Random Talker - Different (talker changed from previous target).

In uninterrupted trials, performance was consistently best in the Same Talker condition, intermediate in the Random Talker - Same condition, and worst in the Random Talker - Different condition (in the left panel of Fig. 6, filled blue bars are tallest and striped bars are shortest at each syllable position). In interrupted trials, performance in the Same Talker condition was generally best for all syllable positions (with performance roughly equal to Random Talker - Same performance on syllable 3). For syllables 3-5, after the interruption, performance was better for Random Talker – Same than for Random Talker – Different; however, for syllable 2, performance was the same whether the talker was the same or differed from the talker on the first syllable.

Linear mixed effect models determined whether or not there were significant differences in performance depending on talker continuity. Models were fitted separately for uninterrupted and interrupted conditions, with fixed effects of talker conditions (Same Talker, Random Talker - Same, Random Talker - Different) and syllable positions (2-5), and a random effect of subjects.

For uninterrupted trials, there were significant main effects of syllable position (*F_3_*,*_473_* = *3.122*, *p* = *0.026*) and talker condition (*F_2_*,*_473_* = *11.495*, *p* < *0.001*), but no significant interaction between them. Post hoc pairwise comparisons showed that performance on target syllable 3 was lower than target syllable 5 (*p* = *0.037*), with no significant differences between other syllable positions. Performance in the Random Talker - Different condition was significantly worse than for either Same Talker (*p* < *0.001*) or Random Talker - Same conditions (*p* = *0.004*), but there was no significant difference between Same Talker and Random Talker - Same condition.

For interrupted trials, our results showed significant main effects of syllable position (*F_3_*,*_473_* = *27.337*, *p* < *0.001*) and talker condition (*F_2_*,*_473_* = *12.628*, *p* < *0.001*), with no significant interaction between them. Post hoc pairwise comparisons showed that performance for syllable 3 was significantly worse performed than target syllable 2, target syllable 4, and syllable 5 (*p* < *0.001* for all). Performance for syllable 5 was also significantly better than for syllables 2 (*p* = *0.010*) and 4 (*p* = *0.003*). Performance for Random Talker - Different was significantly worse than performance for both both Same Talker (*p* < *0.001*) and Random Talker - Same (*p* = *0.003*) conditions, but there was no significant difference between Same Talker and Random Talker - Same conditions.

### B. Discussion

Experiment RT investigated how discontinuity in a task-irrelevant feature (talker) impacts syllable recall in an interrupted spatial selective attention task and how internally generated discontinuities (these random talker changes) interact with an external, salient interrupter. We specifically asked whether discontinuity interferes with streaming and the buildup of spatial attention and thereby increases the impact of interruption.

Randomizing the talker within each stream impaired target recall, demonstrating that talker changes undermine perceptual continuity and weaken spatial selective attention, even though listeners were instructed to focus on spatial cues and ignore talkers. As in earlier experiments, the interrupter produced a large cost on recalling the third target syllable; however, this cost was insignificant on later syllables. Sequential analyses showed that not only was performance better for syllables when the previous syllable was correctly reported, but also when the talker was the same as for the previous target syllable. Together, these results suggest that talker discontinuities behave as internal, unpredictable interruptions that fragment streaming and prevent the buildup of attentional focus over time.

#### 1. Talker discontinuities disrupt attention and streaming

In uninterrupted trials, performance was consistently lower in Random Talker than Same Talker conditions, producing a significant Continuous Talker Benefit. This continuous talker benefit reflects the contribution of automatic spectrotemporal grouping working on top of the attended spatial cue. The sequential analysis reveals why: in the Random Talker condition, performance is worse when the talker changes from syllable to syllable (Random Talker – Different) than when the talker remains the same (Random Talker – Same). Each talker change acts as its own miniature disruption, resetting streaming, thereby interfering with spatial selective attention. Thus, unlike in Experiment CT, where streaming can build steadily in uninterrupted trials, nominally “uninterrupted” Random Talker trials contain repeated resets from talker switches.

In uninterrupted trials, the Continuous Talker Benefit increases from syllable to syllable, suggesting there is a buildup of streaming when the talker doesn’t change; random talker changes break this buildup. This buildup parallels previous reports of a buildup of spatial attention when a target stream comes from the same location from syllable to syllable (Best et al., 2008). However, any kind of interruption, whether from a new event or from a talker switch, interferes with this buildup. In interrupted trials, the external interruption reliably disrupts attention to syllable 3 and also resets streaming between syllables 2 and 3. When streaming is already broken by random talker switches, the interrupter still involuntarily grabs attention to syllable 3, but since streaming is already disrupted, the interrupter does not have a consistent additional effect on syllables 4 and 5.

Many past studies show that there are perceptual costs when processing a speech sequence containing talker switches (Mullennix et al., 1989; H. C. Nusbaum & Morin, 1989). While traditional accounts of this cost argue there is a need for “talker normalization” to correctly interpret acoustically different sounds as the same utterance (e.g., H. Nusbaum & Magnuson, 1997), more recent accounts recognize that abrupt voice changes or discontinuities can also disrupt attention and streaming (e.g., Lim et al., 2021; see Luthra, 2024 for review). Here, we see that the impacts of talker discontinuity are not additive with effects of an external interrupter; instead, both types of disruption break streaming and interfere with attention.

#### 2. Sequential dependencies reveal that talker discontinuities fragment attentional buildup

When the talker remained the same across two adjacent targets—including within the Random Talker condition—performance was substantially better than when the talker changed. Thus, brief stretches of continuity promoted streaming and helped sustain selective attention. Conversely, when the talker switched, the performance was worse, consistent with talker changes causing repeated resets of attentional focus.

The effect of an external interrupter interacted with talker randomization. When an interrupter broke streaming between syllables 2 and 3, performance on syllable 3 depended strongly on whether or not the previous syllable was spoken by the same talker. This improvement due to talker continuity with the previous syllable for syllable 3 was much larger than for other syllable positions.

Together, these results show that talker discontinuities reorient attention much like a salient external interrupter: they break the perceptual continuity of the target stream, reset the buildup of selective attention, and weaken the influence of prior attentional state on subsequent attentional performance. Unlike the single external interrupter, however, random talker switches can produce multiple unpredictable disruptions within a trial. As a result, in the Random Talker condition listeners experience short sequences of syllables that stream together automatically when the talker repeats, separated by random resets. This fragmentation undermines task performance by repeatedly disrupting top-down attention.

#### 3. Conclusions from Experiment RT

Even task-irrelevant acoustic features, when unstable, impose a substantial cost on selective attention. Syllables from the same speaker stream together automatically; when the talker switches unpredictably, the stream itself becomes unstable, making it difficult to maintain top-down spatial focus. Thus, in Experiment RT, the internal discontinuities in the target stream repeatedly reset streaming and undermine attentional focus.

## V. Differences Across Experiments

In Experiment RT, participants showed better performance in interrupted than uninterrupted trials on the first target syllable in Same Talker condition. As noted in Section IV.A.1 and shown in the supplemental materials, we confirmed that the effect on the first syllable was consistent, occurring for the majority of our subjects. This was a surprise. Along with the current Experiment CT, eleven already published experiments tested different versions of the Same Talker condition (Liang et al., 2022, 2025). In none of these experiments was performance worse for any syllable in the interrupted condition compared to the corresponding uninterrupted condition.

### Same Talker performance differs across Experiments CT and RT

To understand why interruption caused performance to be better for the first syllable in Same Talker trials of Experiment RT, we explored whether performance was worse than we typically find on uninterrupted trials, or better on interrupted trials. To this end, we compared performance in the Same Talker conditions in Experiments CT and RT (see Fig. 7). Though we cannot completely rule out other factors that may have differed across the experiments, this comparison of Same Talker results still provides valuable information:

1. In uninterrupted trials, performance was worse in Experiment RT than CT, however, the difference in syllables 1-3 was significantly bigger than that for syllables 4 and 5 (the blue line with open diamonds falls below the blue line with solid circles in Fig. 7, but the distance between the two blue lines is bigger for syllables 1-3 than syllables 4 and 5); and
2. In interrupted trials, performance was similar across the two experiments (dashed red and yellow lines are similar).

**Fig. 7.**
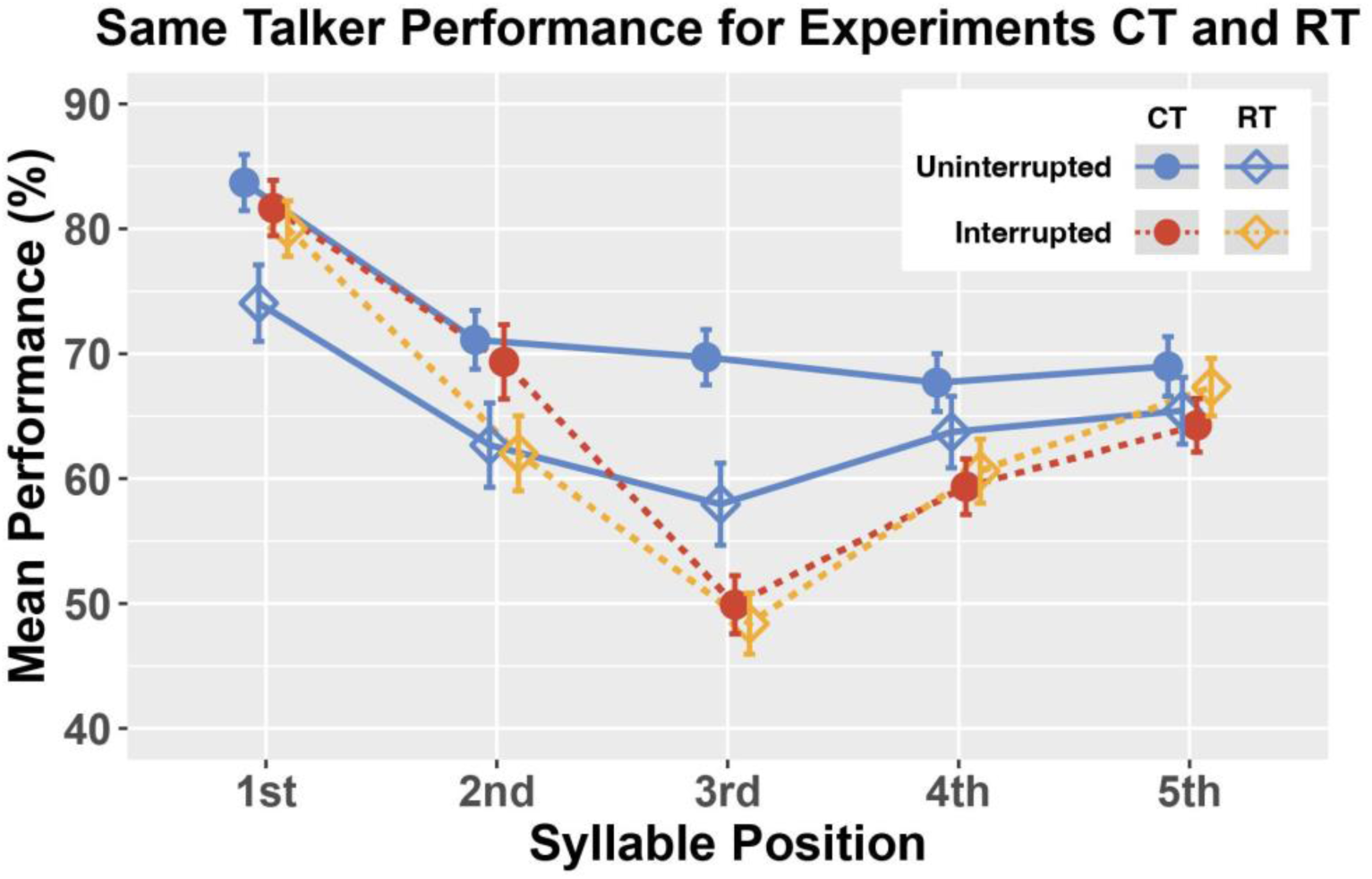
Comparison between syllable recall performance in Experiment CT and RT with the same talker (data replotted from Figures 2 and 4).

Thus, the smaller interruption effect on Targets 1, 2, and 3 in Experiment RT was not because performance was better for interrupted trials; instead, it was due to worse performance on uninterrupted trials.

### Repeating syllable pairs leads to better recall performance in Same Talker conditions

We wondered whether differences in the constraints we applied to the neighboring syllable transitions in the two experiments helps explain differences in the Same Talker condition for Experiments CT and RT. In Experiment CT, syllables 1-4 could match one or both neighboring syllables; however, syllable 5 was guaranteed to differ from syllable 4. In Experiment RT, syllables 1-3 never were the same as any neighboring syllables, but syllables 4 and 5 could match each other.

We explored this by breaking down performance for each syllable, depending upon whether or not a syllable matched one of its neighbors (note: because we are considering only the Same Talker condition, when a syllable matched its neighbor, two matching speech syllables were actually the identical speech token). Figure 8 shows that this has a significant impact on recall: performance was always better when a syllable matched a neighboring syllable than when it differed from its neighbors (for each syllable in each experiment, striped bars, when they are present, are always taller than the corresponding solid bars in Fig. 8).

**Fig. 8.**
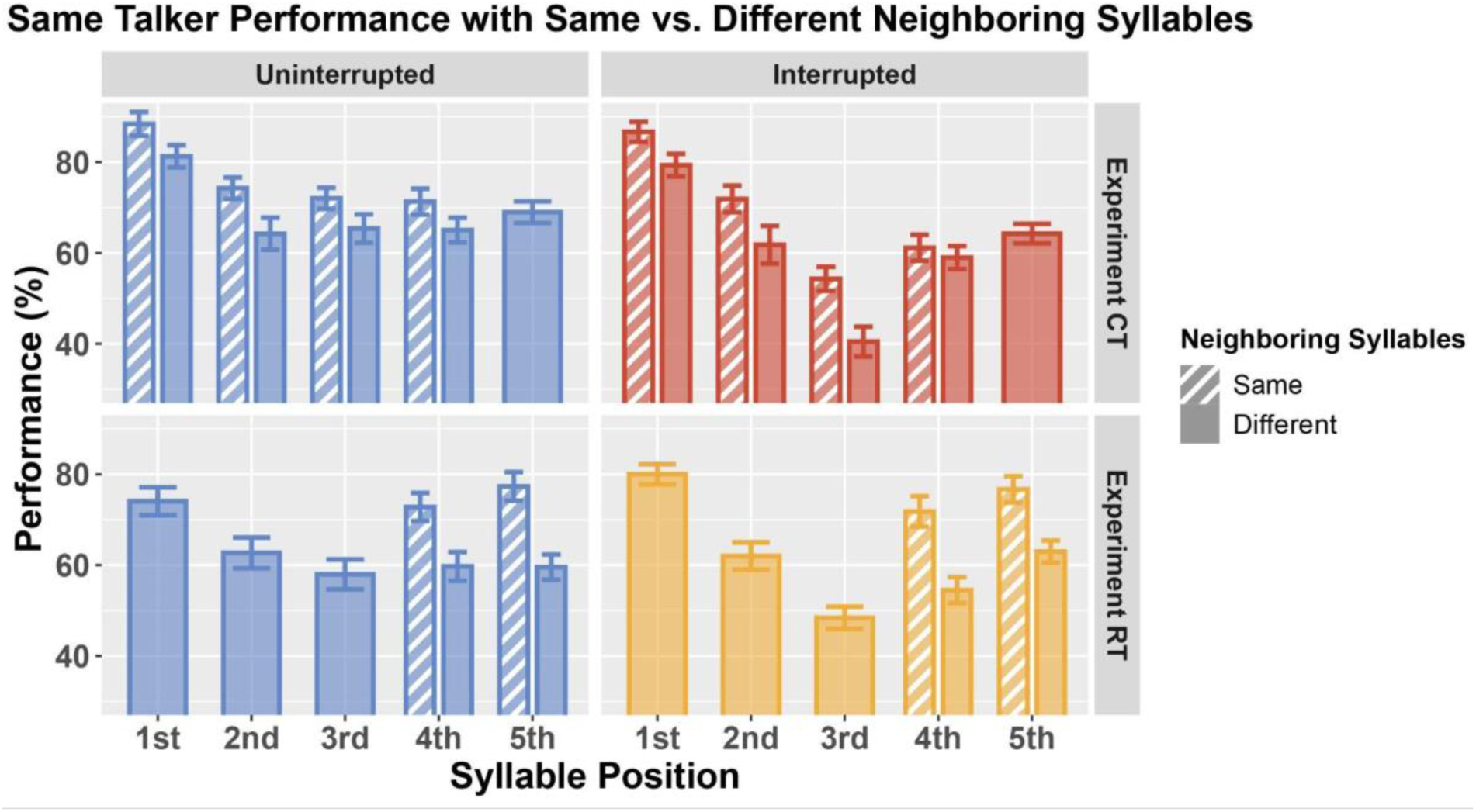
Performance with the same vs. different neighboring syllables, filtered for Same Talker condition.

This analysis demonstrates that recalling two identical syllables appearing in a row is easier than recalling two different syllables. Repeating syllables may be heard as a single unit (a repeating duplet), thereby reducing memory load and improving recall compared to having to store and recall two different syllables. With this in mind, the memory load demands in Experiment RT exceed the load in all of the similar, previously published experiments, where repetition of syllables could occur for every syllable position (Liang et al., 2022, 2025). This high load increases task difficulty overall.

In Experiment CT, the difference in percent correct performance when a syllable repeats a neighbor versus when it differs from its neighbors is consistently less than 10%. In Experiment RT, this difference is larger, generally greater than 10%. This likely reflects differences in the positions that can repeat in the two experiments. In Experiment RT, the repeats only can occur for the last two syllables, while repeats in Experiment CT can occur across the first four syllables. Our results show that the rare, repeating syllables in the final two positions produce especially large benefits of repetition both for uninterrupted and interrupted trials.

We propose two possible, related explanations for this extra-large effect of syllable repetition in Experiment RT: 1) because only the last two syllables ever repeat, this duplet is especially salient, improving recall, and 2) when repeated syllables occur in the last two positions of a sequence, the recency effect in sequential recall is especially effective, boosting performance for the final duplet, not just for the last syllable.

For non-repeating syllables (filled bars in Fig. 8), performance for uninterrupted trials tended to be better in Experiment CT than in Experiment RT (blue bars are higher in the top left panel than the bottom left panel of Fig. 8). This is consistent with load being overall higher in Experiment RT; fewer repeating syllables makes the task harder overall.

Combining these observations:

1. Four of the five syllables in Experiment CT can repeat while only two of the syllables in Experiment RT can match their neighbor, which explains why recall is worse overall in Experiment RT than CT—even on non-repeating syllables.
2. The first few syllables can repeat in Experiment CT, but not in Experiment RT. This further enhances performance on the first few syllables in Experiment CT compared to Experiment RT.
3. The large recall benefit for repetition of the final two syllables improves average performance on these syllables in Experiment RT compared to in Experiment CT, reducing the overall difference in performance for the final two syllables for Same Talker uninterrupted trials.

Together, these effects can account for why Same Talker uninterrupted performance is better overall in Experiment CT, but especially for the first three syllables.

### Interruptions alter recall task demands

Differences in syllable repetition constraints account for why performance for Same Talker uninterrupted trials is better in Experiment CT than in RT (blue open diamonds fall below blue filled circles in Fig. 7). However, performance in interrupted trials is quite similar across the two experiments (red and yellow dashed lines are similar in Fig. 7). One might then assume that repetition has different effects in interrupted trials compared to uninterrupted trials. However, that is not the case. Repetition helps performance nearly the same amount in both uninterrupted and interrupted trials of each experiment (the corresponding left and right panels of Fig. 8 show similar benefits of repetition—differences between dashed and solid bars—at each syllable position). Why, then, do the low number of repeating syllables in Experiment RT not lead to worse overall performance on interrupted trials than in Experiment CT?

We propose that interruption itself can influence task demands, and thus performance, in two distinct ways. First, an interrupter disrupts top-down attention. As a result of disrupted attention, on average fewer target syllables get stored in working memory, reducing memory load overall. Second, when an interrupter breaks perceptual streaming, listeners store the first two syllables as a separate stream, distinct from the final three syllables, rather than as part of a single sequence of five syllables. Such “chunking” of early syllables, separate from the final three, should allow participants to recall these syllables more reliably (Miller, 1956; Ryan, 1969). The idea that interruptions cause syllables 1-2 to be stored and recalled separate from syllables 3-5 gains credence from another piece of evidence. Experiment RT also showed that the benefit of talker continuity is much larger for the *first two* syllables in interrupted trials compared to uninterrupted trials, but is similar for subsequent syllables (in Fig. 4c, for the first two syllables the yellow line is above the blue line, even though the lines lie on top of one another for subsequent syllables). This highlights that in Experiment RT, the first two syllables benefit more from talker continuity in interrupted than uninterrupted trials, as if they are stored and recalled as a distinct pair as long as they do not contain talker discontinuities.

These ideas suggest that interruption can reduce memory load even as it disrupts attention. While there is room for these effects to improve performance in Experiment RT, where memory load is high, they may have limited impact in Experiment CT, where load is already relatively low and patterns of performance more directly reflect interactions between streaming and top-down attentional focus, disruption, and reorientation, not memory capacity.

Together, these effects can explain why performance in Same Talker interrupted trials is similar across our two experiments, even though it differs for uninterrupted trials. On uninterrupted trials, differences in syllable repetition constraints dominate, causing performance to be lower overall in Experiment RT than CT, but especially on the first few syllables. Interruptions reduce load overall, since fewer syllables get into working memory and because the first two syllables are stored and recalled separately from syllables after the interruption. The latter effect is particularly important in counteracting the fact that the first two syllables never repeat in Experiment RT. The effects of interruption on serial recall have a large impact when load is otherwise high, as in Experiment RT, compared to conditions where memory demands are relatively low and not the dominant factor determining performance, as in Experiment CT.

### Interim conclusions, caveats, and future work

A direct comparison of Same Talker trials common to both experiments revealed a large influence of syllable token repetition. Recall is better when a syllable token repeats across neighboring syllables compared to when two different tokens are presented. Because syllables in different positions could repeat (or not) in the two experiments, we found different patterns of performance across experiments. In addition, the relatively low occurrence of repetitions in Experiment RT increased memory load overall, reducing performance.

Even though interruption interferes with top-down attention and hurts performance in general, it also reduces memory load and breaks streaming, allowing syllables before the interruption to be stored and recalled as a separate stream. These effects of interruption offset the costs of increased memory load in Experiment RT, so that interrupted performance was more similar to that in Experiment CT.

Many other factors undoubtedly also contribute to the observed differences across Experiments CT and RT. The subject groups in the two experiments differed, which could contribute to differences in overall performance. In addition, these Same Talker trials we focus on in this section are randomly intermingled with different conditions in the two experiments. In Experiment CT, these other trials (Different Talker condition) were even easier than Same Talker trials; in Experiment RT, these other trials (Random Talker condition) were harder. Thus, the overall task difficulty is greater in Experiment RT than in Experiment CT, which could contribute to fatigue and differences in performance. Yet, these two factors should impact all syllable positions and trial types similarly, and thus do not account for the differences in patterns of performance we observed.

While we propose a number of post hoc explanations for the unexpected differences we found across experiments, future studies should be conducted to test these ideas directly. Specifically, we show that syllable repetition and interruption affect how serial items are chunked and recalled from memory. These may interact with primacy and recency in serial recall, local continuity of talker or other stream features, and sequential dependencies in performance, all of which we observed. Such interactions go beyond what the current results can address but are worthy questions to pursue.

## VI. General Conclusions

In a spatial selective attention task, the task-irrelevant feature of talker identity impacts recall of a target sequence. When two competing streams differ not only in direction, but also in talker, the difference not only helps participants correctly “tag” which syllables are from the target stream when recalling syllables from memory, it allows them to refocus attention rapidly, mitigating the effects of interruption. When the talker of syllables from a particular direction changes randomly, the discontinuity disrupts streaming, breaking spatial attention buildup and interfering with recall. These results support the idea that attention operates on objects: associated spectrotemporal features of syllables, even when not critical to the task, have an obligatory impact on performance.

Provocatively, spatial features, which can be easily deployed to focus top-down attention, may *not* be automatically bound to items in auditory memory. Specifically, when attention is interrupted and two concurrent streams are spoken by the same talker, performance on the syllable occurring right after the interrupter is typically low, as if the disruption of top-down spatial attention “wipes out” spatial information in short-term memory. The largest benefit of talker differences occur for this syllable—demonstrating that talker differences reliably identify which syllable comes from the target.

Sequential effects in recall demonstrate further influences of perceptual streaming and continuity on task performance. In all cases, listeners recall a particular syllable more reliably when they correctly reported the previous syllable. However, these sequential effects are stronger when target and distractor streams are distinct (from different talkers) and continuous (not interrupted by a random sound and not containing talker discontinuities).

Repetition of neighboring syllable tokens has a large influence on serial recall performance. Syllable repetition has a local effect, improving recall of the repeating tokens, as well as a global effect, reducing task difficulty overall.

Interruptions during a spatial selective attention task not only interfere with top-down focus, they break perceptual continuity of the attended stream. This can cause what would otherwise be a continuous target stream to be perceived and stored in separate pieces.

This study builds on previous studies of the effects of interruption on top-down spatial selective attention (Liang et al., 2022, 2025) by exploring how a task-irrelevant feature of talker influences performance during disruption. Results demonstrate that even features that participants are instructed to ignore fundamentally influence the perceptual organization of the auditory scene, impacting how listeners stream and store information and how they recover from unexpected interruptions.

## Ethical Approval

This study was reviewed and approved by the Carnegie Mellon University Institutional Review Board (STUDY2019_00000217).

## Consent to Participate

All participants provided written informed consent for their participation.

## Data Availability Statement

The datasets generated during and/or analyzed during the current study are available in the CMU KiltHub repository, 10.1184/R1/30870710.

## REFERENCES

Bee, M. A., & Micheyl, C. (2008). The cocktail party problem: what is it? How can it be solved? And why should animal behaviorists study it? Journal of Comparative Psychology (Washington, D.C.: 1983), 122(3), 235–251.

Best, V., Ozmeral, E. J., Kopco, N., & Shinn-Cunningham, B. G. (2008). Object continuity enhances selective auditory attention. Proceedings of the National Academy of Sciences of the United States of America, 105(35), 13174–13178.

Bizley, J. K., & Cohen, Y. E. (2013). The what, where and how of auditory-object perception. Nature Reviews. Neuroscience, 14(10), 693–707.

Bonacci, L. M., Bressler, S., & Shinn-Cunningham, B. G. (2020). Nonspatial Features Reduce the Reliance on Sustained Spatial Auditory Attention. Ear and Hearing, 41(6), 1635–1647.

Bressler, S., Masud, S., Bharadwaj, H., & Shinn-Cunningham, B. (2014). Bottom-up influences of voice continuity in focusing selective auditory attention. Psychological Research, 78(3), 349–360.

Bronkhorst, A. W. (2015). The cocktail-party problem revisited: early processing and selection of multi-talker speech. Attention, Perception & Psychophysics, 77(5), 1465–1487.

Cherry, E. C. (1953). Some Experiments on the Recognition of Speech, with One and with Two Ears. The Journal of the Acoustical Society of America, 25(5), 975–979.

Fischer, C., Nolting, C., Schneider, F., Bledowski, C., & Kaiser, J. (2024). Auditory objects in working memory include task-irrelevant features. Scientific Reports, 14(1), 21216.

Gardner, B., & Martin, K. (n.d.). HRTF Measurements of a KEMAR DummyHead Microphone. Retrieved December 5, 2023, from http://www.linux.bucknell.edu/~kozick/elec32007/hrtfdoc.pdf

Huang, N., & Elhilali, M. (2020). Push-pull competition between bottom-up and top-down auditory attention to natural soundscapes. eLife, 9, e52984.

Lee, C.-C., & Middlebrooks, J. C. (2011). Auditory cortex spatial sensitivity sharpens during task performance. Nature Neuroscience, 14(1), 108–114.

Liang, W., Brown, C. A., Noyce, A. L., & Shinn-Cunningham, B. G. (2025). Cat-astrophic update: What makes an interrupter more disruptive? The Journal of the Acoustical Society of America, 158(5), 4048–4058.

Liang, W., Brown, C. A., & Shinn-Cunningham, B. G. (2022). Cat-astrophic effects of sudden interruptions on spatial auditory attention. The Journal of the Acoustical Society of America, 151(5), 3219.

Lim, S.-J., Carter, Y. D., Michelle Njoroge, J., Shinn-Cunningham, B. G., & Perrachione, T. K. (2021). Talker discontinuity disrupts attention to speech: Evidence from EEG and pupillometry. In Brain and Language (Vol. 221, p. 104996). 10.1016/j.bandl.2021.104996

Luthra, S. (2024). Why are listeners hindered by talker variability? Psychonomic Bulletin & Review, 31(1), 104–121.

Michalka, S. W., Kong, L., Rosen, M. L., Shinn-Cunningham, B. G., & Somers, D. C. (2015). Short-Term Memory for Space and Time Flexibly Recruit Complementary Sensory-Biased Frontal Lobe Attention Networks. Neuron, 87(4), 882–892.

Middlebrooks, J. C., & Waters, M. F. (2020). Spatial Mechanisms for Segregation of Competing Sounds, and a Breakdown in Spatial Hearing. Frontiers in Neuroscience, 14, 571095.

Miller, G. A. (1956). The magical number seven plus or minus two: some limits on our capacity for processing information. Psychological Review, 63(2), 81–97.

Milne, A. E., Bianco, R., Poole, K. C., Zhao, S., Oxenham, A. J., Billig, A. J., & Chait, M. (2021). An online headphone screening test based on dichotic pitch. Behavior Research Methods, 53(4), 1551–1562.

Mullennix, J. W., Pisoni, D. B., & Martin, C. S. (1989). Some effects of talker variability on spoken word recognition. The Journal of the Acoustical Society of America, 85(1), 365–378.

Nusbaum, H. C., & Morin, T. M. (1989). Perceptual normalization of talker differences. The Journal of the Acoustical Society of America, 85(S1), S125–S125.

Nusbaum, H., & Magnuson, J. S. (1997). Talker normalization: Phonetic constancy as a cognitive process. ResearchGate.

Pressnitzer, D., Sayles, M., Micheyl, C., & Winter, I. M. (2008). Perceptual organization of sound begins in the auditory periphery. Current Biology: CB, 18(15), 1124–1128.

Qian, Y.-M., Weng, C., Chang, X.-K., Wang, S., & Yu, D. (2018). Past review, current progress, and challenges ahead on the cocktail party problem. Frontiers of Information Technology & Electronic Engineering, 19(1), 40–63.

Ryan, J. (1969). Grouping and short-term memory: different means and patterns of grouping. The Quarterly Journal of Experimental Psychology, 21(2), 137–147.

Zhao, S., Yum, N. W., Benjamin, L., Benhamou, E., Yoneya, M., Furukawa, S., Dick, F., Slaney, M., & Chait, M. (2019). Rapid ocular responses are modulated by bottom-up-driven auditory salience. The Journal of Neuroscience: The Official Journal of the Society for Neuroscience, 39(39), 7703–7714.

