## supplemental_material for "Distinct talkers combat catastrophic failures of spatial attention due to interruption"

***Experiment RT: Majority performed syllable 1 better in interrupted than uninterrupted trials***

To investigate the better performance for syllable 1 in interrupted than uninterrupted trials in Same Talker condition in Experiment RT, we looked at individual performance. Analysis confirmed that the majority of subjects performed better for syllable 1 in interrupted trials than uninterrupted trials (Fig. S1). Each line connects an individual's performance in uninterrupted and interrupted trials. The color of the line is determined by which condition a particular individual performed better: blue lines indicate better performance in uninterrupted trials, yellow indicate better performance in interrupted trials, gray indicates equal performance in uninterrupted and interrupted trials.

The majority of the lines were yellow for the first syllable (25 of 44 participants), indicating the majority of the participants performed better in interrupted trials than uninterrupted trials. For all subsequent syllables, as in previous, similar experiments for all syllable positions, the majority of participants performed better in uninterrupted trials than interrupted trials (majority blue lines).

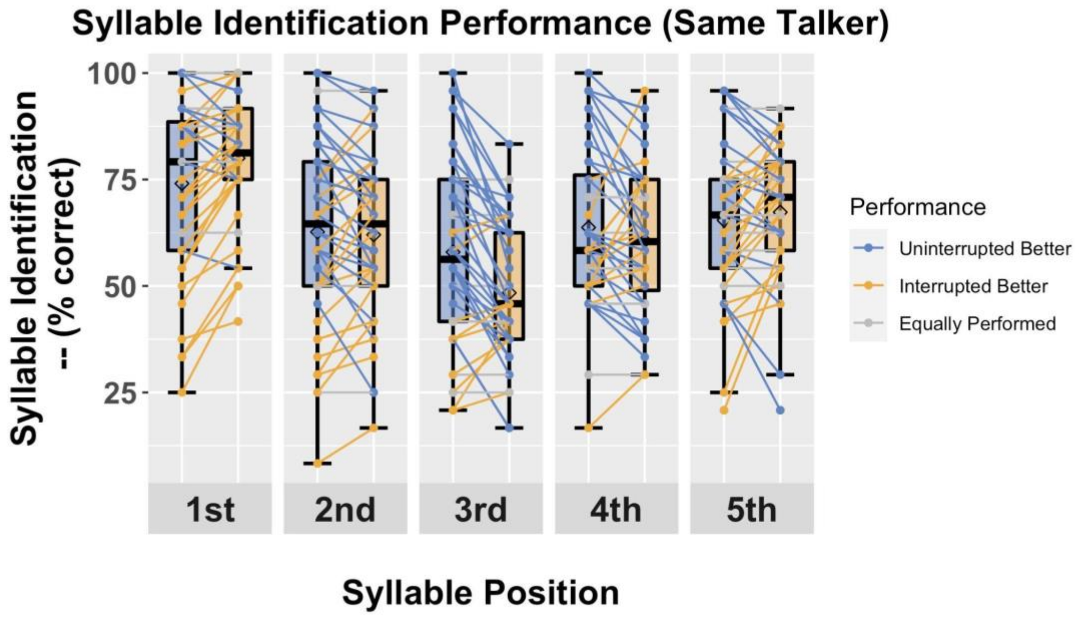


*Fig. S1 Syllable recall performance with the same talker for target and distractor streams. Lines connecting each individual's performance in uninterrupted and interrupted trials. Blue line means that the individual performed better in uninterrupted trials at that syllable position, yellow line means the individual performed better in interrupted trials, gray line means equal performance in uninterrupted and interrupted conditions. From observation, the majority of the subjects performed better in interrupted trials for the first syllable position.*
